# Insulin enhances presynaptic glutamate release via opioid receptor-mediated disinhibition

**DOI:** 10.1101/517797

**Authors:** Tracy L. Fetterly, Max F. Oginsky, Allison M. Nieto, Yanaira Alonso-Caraballo, Zuleirys Santana-Rodriguez, Carrie R. Ferrario

## Abstract

Insulin influences activity in brain centers that mediate reward and motivation in humans. However, nothing is known about how insulin influences excitatory transmission in regions like the nucleus accumbens (NAc), which governs motivational processes in the adult brain. Further, insulin dysregulation that accompanies obesity is linked to cognitive decline, depression, anxiety, and aberrant motivation that also rely on NAc excitatory transmission. Using a combination of whole-cell patch clamp and biochemical approaches we determined how insulin affects NAc glutamatergic transmission. We show that insulin receptor activation increases presynaptic glutamate release via a previously unidentified form of opioid receptor-mediated disinhibition. In contrast, activation of IGF receptors by insulin decreases presynaptic glutamate release in adult male rats. Furthermore, obesity results in a loss of insulin receptor-mediated increases and a reduction in NAc insulin receptor surface expression, while preserving reductions in transmission mediated by IGRFs. These results provide the first insights into how insulin influences excitatory transmission in the adult brain, they provide foundational information about opioid-mediated regulation of NAc glutamatergic transmission, and have broad implications for the regulation of motivation and reward related processes by peripheral hormones.

## I. INTRODUCTION

Recent studies in humans and rodents suggest that insulin may enhance cognition and decision-making processes that influence reward seeking (Reger et al., 2008; Freiherr et al., 2013). However, the mechanisms by which insulin affects neural function in the adult brain are poorly understood (Biessels and Reagan, 2015; Ferrario and Reagan, 2017). A few studies primarily in cortical and hippocampal neurons have shown that insulin influences excitatory transmission via pre-synaptic mechanisms that reduce glutamate release, as well as post-synaptic mechanisms that affect AMPAR trafficking (Beattie et al., 2000; Man et al., 2000; Passafaro et al., 2001; Huang et al., 2004; Labouebe et al., 2013; Liu et al., 2013). However, these studies were conducted in cultured neurons or juvenile rodents (18-30 days old). Thus, very little is known about effects of insulin on excitatory transmission in the adult brain.

Glutamatergic transmission within the nucleus accumbens (NAc) mediates many aspects of motivation and decision-making in response to food, sex, and drugs of abuse, as well as to environmental stimuli paired with these rewards. For example, food- and drug-seeking behaviors rely on activation of the NAc (Di Ciano et al., 2001; Kalivas, 2009; Wolf, 2016), and repeated exposure to drugs of abuse or palatable foods enhances NAc excitatory transmission that underlies food- and drug-seeking behaviors (Oginsky et al., 2016; Wolf, 2016; Dong et al., 2017; Alonso-Caraballo et al., 2018; Derman and Ferrario, 2018; Alonso-Caraballo et al., In Press). Thus, identifying neural mechanisms that regulate NAc excitatory transmission is fundamental to understanding the neurobiology of normal and aberrant motivation.

Here, we used whole-cell patch clamp recordings in adult rat brain slices to determine how insulin affects excitatory transmission onto NAc medium spiny neurons (MSNs) and the mechanisms involved. Importantly, in addition to insulin receptors, insulin-like growth factor receptors (IGFRs) are also expressed in the NAc and can be activated by moderate to high concentrations of insulin (Unger et al., 1989; Schumacher et al., 1991). Thus, a wide range of insulin concentrations were examined here, and the contribution of insulin receptor activation vs. IGFR activation to insulins effects were determined. In addition, given that obesity is associated with insulin dysregulation, altered NAc excitatory transmission, cognitive deficits and some psychiatric diseases (Biessels and Reagan, 2015; Kullmann et al., 2016; Stoeckel et al., 2016), we also determined how high-fat diet induced obesity alters insulins ability to influence NAc excitatory transmission.

We found that insulin receptor and IGFR activation have opposing effects on excitatory transmission in the NAc, with insulin receptor activation increasing and IGFR activation decreasing presynaptic glutamate release. Furthermore, insulin-induced increases in glutamate release occurred through a previously unidentified opioid receptor-mediated form of disinhibition. Finally, diet-induced obesity resulted in a loss of insulin-induced increases in NAc excitatory transmission and a reduction in NAc insulin receptor surface expression. Together, these data reveal novel roles for insulin in the regulation of NAc excitatory transmission, provide new insights into opioid-mediated regulation of NAc glutamatergic transmission, and have implications for endogenous and exogenous insulin in modulating motivation and reward.

## II. METHODS

### Animals

Male Sprague-Dawley rats were purchased from Envigo (Indianapolis, IN), pair-housed (reverse light-dark; 12/12) with free access to food and water unless otherwise stated (70-80 days old). All procedures were approved by the UM Institutional Animal Care and Use Committee (IACUC). Additional information about housing conditions and procedures can be found here: https://sites.google.com/a/umich.edu/ferrariolab-public-protocols/

### Electrophysiology

Whole-cell patch-clamp recordings of MSNs in the NAc core were conducted as previously described (Ferrario et al., 2011; Oginsky et al., 2016). Rats were anesthetized with chloral hydrate (400 mg/kg, i.p.), brains were rapidly removed and placed in ice-cold oxygenated (95% O2-5% CO2) aCSF containing (in mM): 125 NaCl, 25 NaHCO3, 12.5 glucose, 1.25 NaH2PO4, 3.5 KCl, 1 L-ascorbic acid, 0.5 CaCl2, 3 MgCl2, 295-305 mOsm, pH 7.4. Coronal slices (300 *µm*) containing the NAc were made using a vibratory microtome (Leica Biosystems, Buffalo Grove, IL, USA) and allowed to rest in oxygenated aCSF (40 min). For the recording aCSF (2 ml/min), CaCl2 was increased to 2.5 mM and MgCl2 was decreased to 1 mM. Patch pipettes were pulled from 1.5 mm borosilicate glass capillaries (WPI, Sarasota, FL; 37 MΩ resistance) and filled with a solution containing (in mM): 140 CsCl, 10 HEPES, 2 MgCl2, 5 Na+-ATP, 0.6 Na+-GTP, 2 QX314, pH 7.3, 285 mOsm. All recordings were conducted in the presence of picrotoxin (50 *µ* M).

Evoked EPSCs (eEPSCs) were elicited by local stimulation (0.05 to 0.30 mA square pulses, 0.3 ms, delivered every 20 s) using a bipolar electrode placed approximately 300 microns lateral to recorded neurons. The minimum amount of current needed to elicit a synaptic response with <15% variability in amplitude was used. If > 0.30 mA was required, the neuron was discarded. eEP-SCs were recorded at −70 mV. Baseline responses were established (10 min) followed by a bath application of insulin in the presence or absence of antagonists (10 min). mEPSCs were recorded in the presence of tetrodotoxin (1 *µM*) at a holding potential of −65mV. To validate paired-pulse facilitation procedures, eEPSCs were measured across a range of inter-pulse intervals (50, 75, 100, 200 and 400 ms; 6-8 pulses per interval) in the same cell. The probability of glutamate release was determined by dividing the averaged amplitude of the 2nd peak by the averaged amplitude of the 1st peak (i.e., paired pulse ratio).

Recorded signals were amplified with a Multiclamp 700B (Molecular Devices, Union City, CA), digitized at 20 kHz and filtered at 2 kHz and collected with Clampex 10.4 data acquisition software (Molecular Devices). All drugs were bath applied for 10 min (Sigma Aldrich: insulin [91077C], phaclofen [114012-12-3], (-)- naloxone [51481-60-8], bestatin [65391-42-6], thiorphan [76721-89-6]), HNMPA and HNMPA-(AM)3 (Santa Cruz Biotechnology; sc-205714, sc-221730), picropodophyllotoxin (PPP, Tocris cat# 2956), (+)-naloxone was provided by Kenner C. Rice (Drug Design and Synthesis Section, NIDA IRP). In our initial studies, insulin concentrations ranging from 1 to 500 nM were used. This was done in part to facilitate comparison to effects of insulin in other brain regions including the VTA where concentrations of 100 and 500 nM have been used (Labouebe et al., 2013; Liu et al., 2013) and to avoid missing effects by examining just one concentration. Furthermore, while physiological concentrations of insulin are thought to be relatively low (10-30 nM), how these levels may be affected by diet-induced obesity and/or the diabetic state is not understood and therefore levels could be much higher (see Ferrario and Reagan, 2018 for additional discussion).

### Single Cell RT-PCR and Identification of D1- and D2-type MSNs

Single-cell RT-PCR was conducted on cell contents taken from MSNs after whole-cell recordings to identify D1-and D2-type MSNs. The first-strand cDNA synthesis was performed using the Superscript III First-Strand Synthesis System for RT-PCR (Life Technologies, Grand Island, NY) per the manufacturer’s instructions. The reverse transcription product was kept at 20°*C* until PCR was performed. PCR primers used: prodynorphin forward: 5-GCCTAGGAGTGGAGTGTTCG, reverse: 5-GGGATAGAGCAGTTGGGCTG; proenkephalin forward: 5-ATGCCATGCCATCGGGAAG, reverse: 5-CAGGACCAGCAGGGACAATC. PCR product lengths were >100 bp so as to not confuse them with primer dimers. Four *µL* (prodynorphin) or six *µL* (proenkephalin) of reverse transcription product were loaded into an Eppendorf tube with PCR solution containing 10 *µL* of 5 green GoTaq flexi buffer, 2 *µL* MgCl2, 1*µL* of 10 mM dNTP mix, 1 *µL* of 10 mM forward and reverse primers, 0.25 *µL* of GoTaq polymerase (Promega), and brought up to a final volume of 50 *µL* with nuclease-free water. The thermal cycling program was set to the initial denaturation for 5 min at 95°*C* for one cycle. The denaturation, annealing, extension cycles were done at 95°*C* for 1 min, 58°*C* (Enk) and 65°*C* (Dyn) for 1 min, and 72°*C* for 1 min, respectively, for 45 cycles. A final extension cycle was done at 72°*C*for 5 min. Four *µL* of the PCR reaction were placed into a second PCR tube with the same solution as before and the same cycling protocol was performed. Twenty *µL* from the second PCR reaction was run on a 2% agarose gel containing ethidium bromide. Gels were imaged using UVP GelDoc-It2 imager (UVP, Upland, CA). D1- or D2-type MSNs were defined by the presence of a PCR product band for either prodynorphin or proenkephalin, respectively.

### High-fat Diet-induced Obesity

Rats were given free access to 60% high-fat diet (Open Source Diets D12492) in the home cage for a total of 8 weeks. Controls had free access to standard lab chow throughout (Lab Diet 5001, 13% fat). Weight was measured twice each week. In addition, after 7 weeks of high-fat or control diet, body composition was determined by NMR (Minispec LF90II, Bruker Optics), and fasted blood samples (16 hrs) were collected and used to determine plasma insulin levels. Blood samples were collected into tubes containing EDTA (1.6 mg/mL, Sarstedt) and plasma was then isolated by centrifugation (1000 x G, 4 °*C*, 10 min), and stored (−20 °*C*) for subsequent analysis as previously described (Vollbrecht et al., 2015). Plasma insulin levels were determined by double-antibody radioimmunoassay using a 125I-Human insulin tracer (Linco Research St. Charles, MO), a rat insulin standard (Novo Nordisk, Plainsboro, NJ), a guinea pig anti-rat insulin first antibody (Linco Research), and a sheep anti-guinea pig gamma globulin-PEG second antibody (Michigan Diabetes Research Core). Blood collection and NMR were conducted at week 7 in order to avoid additional stress during the week of slice preparation or NAc tissue collection (week 8). Food was removed from the cage 1-2 hours before slice preparation or NAc tissue collection.

### Purification of Surface (bound) Proteins, and Western Blotting

NAc tissue was biotinylated and NeutrAvidin isolation of biotinylated (surface) proteins were conducted as previous described (Ferrario et al., 2011). For these experiments, verification studies were done to determine optimal pull down procedures and the amount of material to be loaded per lane. Briefly, bilateral NAc tissue (containing core and shell) from each rat was dissected and chopped (400 *µm*; McIllwain tissue chopper; the Vibratome Company, OFallon, MO). NAc tissue was added to ice-cold aCSF containing 1 mM sulfo-NHS-S-S-Biotin (Thermo Scientific, Rockford, IL) and incubated with gentle agitation (30 min, 4°*C*). This reaction was quenched by the addition of glycine (100 mM, 10 min, 4°*C*), tissue was pelleted, and re-suspended in ice-cold lysis buffer (in mM: 25 HEPES; 500 NaCl, 2 EDTA, 1 phenylmethyl sulfonyl fluoride, 20 NaF, 1:100 protease inhibitor cocktail set I [Calbiochem, San Diego, CA], and 0.1% Nonidet P-40 [v/v]; pH 7.4), sonicated and stored at −80°*C* for subsequent use.

Procedures to purify biotinylated (i.e., surface) proteins were adapted from Thermo Scientific product instructions and all steps were conducted on ice or at 4°*C* unless otherwise noted. Protein concentrations were determined by Pierce BCA assay. 100 *µg* of NAc protein was added to high capacity NeutrAvidin agarose beads (Thermo Scientific, Cat #29202) and incubated overnight with end-over-end rotation. Biotinylated proteins bound to NeutrAvidin beads (bound, surface fraction) were isolated from the non-biotinylated (unbound) fraction by centrifugation (3000 RPM, 1 min) and washed (3 times, 1 X PBS). The supernatant (unbound) was collected and fresh beads were added for a second overnight incubation and isolation of surface proteins as above. The bound fractions were combined in a total of 70 *µL* of Laemmli sample treatment buffer containing DTT (100 mM), and heated at 97°*C* for 3 min to release the biotinylated proteins from the beads. The bound samples were then spun at 10,000 RPM for 5 min on a centrifugal filter unit (0.45 mm, #UFC30HV00, Millipore, Billerica, MA) to remove the NeutrAvidin beads from the solution. The samples were then stored at −20°*C* until used for Western Blotting (Chow N=10, High-fat N=10).

For Western blotting, bound fractions (surface protein) or whole cell lysates (total protein) were heated (70°*C*, 10 min), loaded into gels (20 *µg* whole cell lysate, 20 *µL* bound fraction) and electrophoresed under reducing conditions. Proteins were transferred onto PVDF membranes (Millipore, Cat # IPVH 00010), membranes were rinsed, blocked (1hr, RT, 5% [w/v] nonfat dry milk in TBS-Tween 20 [TBS-T; 0.05% Tween 20, v/v]), and incubated overnight with primary antibody to the beta subunit of the insulin receptor (IR; 1:200 in TBS; Santa Cruz S711). To verify that intracellular proteins were not abundant in the bound fraction, the relative expression of tyrosine hydroxylase (TH, 1:30,000 in TBS; Life Technologies, P21962) was determined in the bound and unbound fractions. Membranes were then washed in TBS-T, incubated with HRP-conjugated secondary (Invitrogen, Carlsbad, CA; 1hr, RT), washed, and immersed in chemiluminescence detecting substrate (GE Health-care, Piscataway, NJ). Images were acquired on film and Ponceau S (Sigma-Aldrich) was used to determine total protein in each lane. Bands of interest were quantified using Image J (NIH).

### Quantification and Statistical Analysis

eEPSCs were analyzed with Clampfit 10.4 (Molecular Devices), mEPSCs were analyzed using Mini Analysis program 6.0.4 (Synaptosoft) and verified by hand. No more than 3 cells were included/rat for any given measure to avoid over-representation of one subject. Two-tailed t-tests, one-way or two-way repeated measures ANOVAs, and Sidaks post-hoc multiple comparisons tests were conducted using Prism 6-8 software (GraphPad). Statistical tests used for each data set are stated in the results section below and in brief in the figure captions. Ns are given in the results section, with the number of cells followed by the number of rats used for electrophysiological recordings (e.g., 6,5 = 6 cells from 5 rats). Data in all figures are shown as average *±* SEM.

## III. RESULTS

### A. Insulin bi-directionally influences NAc excitatory transmission

Using established whole-cell patch clamping approaches from adult brain slices (Ferrario et al., 2011; Oginsky et al., 2016), we first determined how bath application of insulin (1-500 nM) affects the amplitude of evoked excitatory postsynaptic currents (eEPSC) in MSNs of the NAc core (Figure 1). We found that 30nM insulin significantly increased eEPSC amplitude (Figure 1A closed circles; Two-way RM ANOVA main effect 30nM: *F*_(1,7)_=10.55, p=0.01; N=7,6), whereas 100 or 500nM insulin produced a significant decrease in amplitude (Figure 1A triangles, diamonds; two-way RM ANOVA main effect 100nM: *F*_(1,4)_=19.56, p=0.01; N=5,4; main effect 500nM: *F*_(1,4)_=43.50, p=0.003; N=5,4). In addition, eEPSC amplitude returned to baseline following insulin washout. Furthermore, eEPSC amplitude was unchanged by 50nM (Figure 1A squares; twoway RM ANOVA main effect 50nM: *F*_(1,3)_>0.000006, p=0.99; N=4,3), 1nM or 10nM insulin (Figure 1B, heptagon, triangle; two-way RM ANOVA main effect 1nM: *F*_(1,5)_=0.093, p=0.77; N=6,4; main effect 10nM: *F*_(1,3)_=1.617, p=0.29; N=4,2). Thus, insulin produces bidirectional and concentration dependent effects on NAc excitatory transmission.

**FIG. 1:**
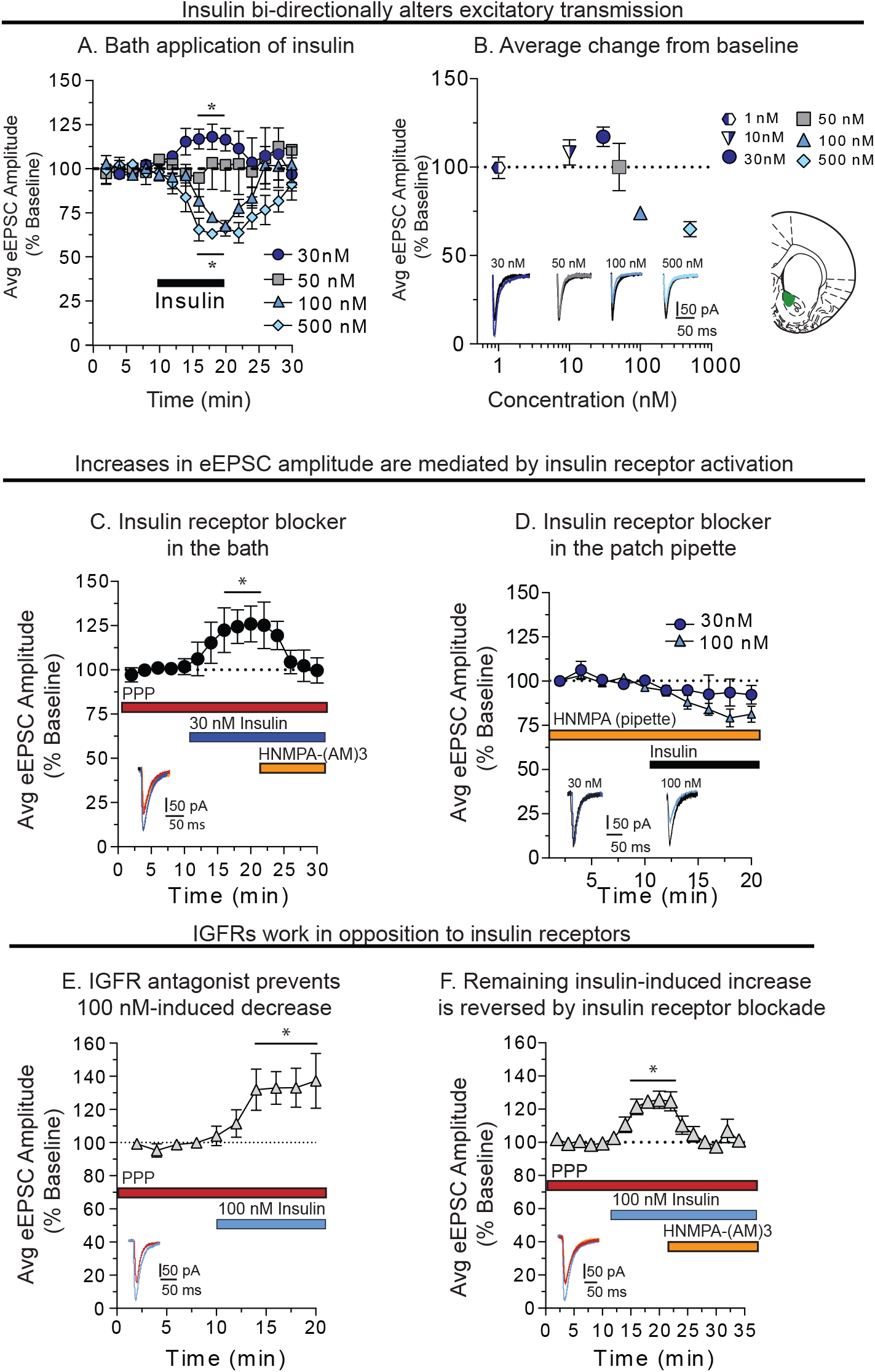
A) Average eEPSC amplitude during baseline, bath application of insulin (black bar) and following washout. B) Summary of average maximum change from baseline following insulin (1-500nM). Recording location within the NAc core is shown at the right. C) Average eEPSC amplitude in the presence of the IGFR-antagonist PPP, before and after 30nM insulin, with and without the membrane permeable insulin receptor inhibitor HNMPA-(AM)3. D) Average eEPSC amplitude before and after insulin (30 and 100nM) with the membrane-impermeable insulin receptor inhibitor HNMPA included in the patch pipette. E) Average eEPSC amplitude before and after 100nM insulin administered in the presence of PPP. F) Average eEPSC amplitude in the presence of PPP, before and after 100nM insulin followed by the addition of HNMPA-(AM)3 to the bath. Statistical differences were determined by two-way RM ANOVA comparing baseline and treatment conditions. ∗= main effect of treatment, p < 0.01.

### B. Insulin receptor and IGFR activation have opposing effects on NAc excitatory transmission

In adult brain, insulin activates insulin receptors and IGFRs (Vigneri et al., 2010). However, because of the different binding affinities of these receptors, low concentrations of insulin (∼ 30 nM) preferentially activate insulin receptors, whereas higher concentrations also activate IGFRs (Schumacher et al., 1991). We therefore hypothesized that increases in excitatory transmission elicited by 30 nM insulin may be mediated by insulin receptors, whereas decreases following 100 nM may be mediated by IGFRs. To test this, we bath applied selective antagonists of the IGFR (picropodophyllotoxin [PPP], 500 nM; Labouebe et al., 2013) or the insulin receptor blocker (HNMPA-(AM)3; 100 *µM*, Saperstein et al., 1989; Mebel et al., 2013) before or after insulin (Figure 1C-F). Additional controls were conducted to assess the effect of these drugs on baseline eEPSC amplitude. We found that PPP increased eEPSC amplitude on its own (data not shown; two-way RM ANOVA main effect PPP: *F*_(1,8)_=24.0, p=0.001; N=5,3). Therefore, PPP was always applied prior to further drug manipulations to allow for a stable baseline to be established. In addition, application of HNMPA-(AM)3 to the bath alone did not alter eEPSC amplitude (data not shown; two-way RM ANOVA main effect HNMPA-(AM)3: *F*_(1,3)_=0.004, p=0.95; N=4,3).

Consistent with our hypothesis, application of 30nM insulin in the presence of the IGFR antagonist PPP resulted in a significant increase in eEPSC amplitude that was revered by the subsequent addition of the membrane permeable insulin receptor blocker, HNMPA-(AM)3 to the bath (Figure 1C; two-way RM ANOVA condition x time interaction: *F*_(8,48)_=2.91, p=0.01; N=7,4). Furthermore, when the membrane-impermeable insulin receptor blocker HNMPA (300 *µM*) was included in the recording pipette 30 nM insulin-induced increases in eEPSC amplitude were also completely blocked (Figure 1D; two-way RM ANOVA *F*_(1,4)_=2.7, p=0.17; N=5,4), while 100nM insulin-induced decreases in eEPSC amplitude were still observed (Figure 1D, triangles; two-way RM ANOVA main effect 100nM insulin: *F*_(1,8)_=19.2, p=0.002; N=9,5). As this manipulation would only prevent activation of insulin receptors within the recorded MSN, these data indicate that increases in excitatory transmission are due to activation of insulin receptors located on MSNs.

When the IGFR antagonist PPP was applied prior to 100 nM insulin, previously observed decreases in eEPSC amplitude were absent (Figure 1E). Furthermore, under this condition insulin produced a significant increase in eEPSC amplitude (Figure 1E; two-way RM ANOVA main effect 100nM + PPP: *F*_(1,5)_=7.55, p=0.04; N=6,4), likely due to activation of insulin receptors (which are not blocked by PPP). To verify this, PPP was included in the bath followed by 100 nM insulin with and without HNMPA-(AM)3. Under these conditions, insulin-induced increases were completely reversed by the insulin receptor blocker (Figure 1F; two-way RM ANOVA main effect of condition: *F*_(2,16)_=12.5, p<0.001; N=9,5). Together, these data demonstrate that insulin receptors and IGFRs work in opposition to enhance and reduce NAc excitatory transmission, respectively.

### C. Identification of D1-type and D2-type MSNs after whole-cell recording

MSNs can be sub-divided by their expression of D1- and D2-like receptors, with each population having dissociable roles in motivated behavior (Yager et al., 2015). Within the NAc, D1-type MSNs project to the substantia nigra and VTA (output nuclei), whereas D2- and D1-MSNs project to the ventral pallidum, which is a relay as well as an output nucleus. Compared to studies in the dorsal portion of the striatum, relatively little is known about potential differences in the regulation of excitatory transmission onto D1-vs and D2-MSNs. Therefore, as a first step towards examining potential differences in insulin’s effects on these two populations, we established single cell RT-PCR approaches following whole-cell patch clamping to classify a subset of recorded neurons as D1- or D2-type MSNs (cells from data in Figs 1, 3, and 5). D2-MSNs were identified by the presence of proenkephalin (pEnk) an absence prodynoprhin (pDyn), whereas D1-MSNs were identified by the opposite pattern (Figure 2A). We also determined that the sensitivity of pEnk primers was lower than that of pDyn primers (compare Figure 2B, C), and that sensitivity of pENK primers can be enhanced by additional amplification (Figure 2D).

**FIG. 2:**
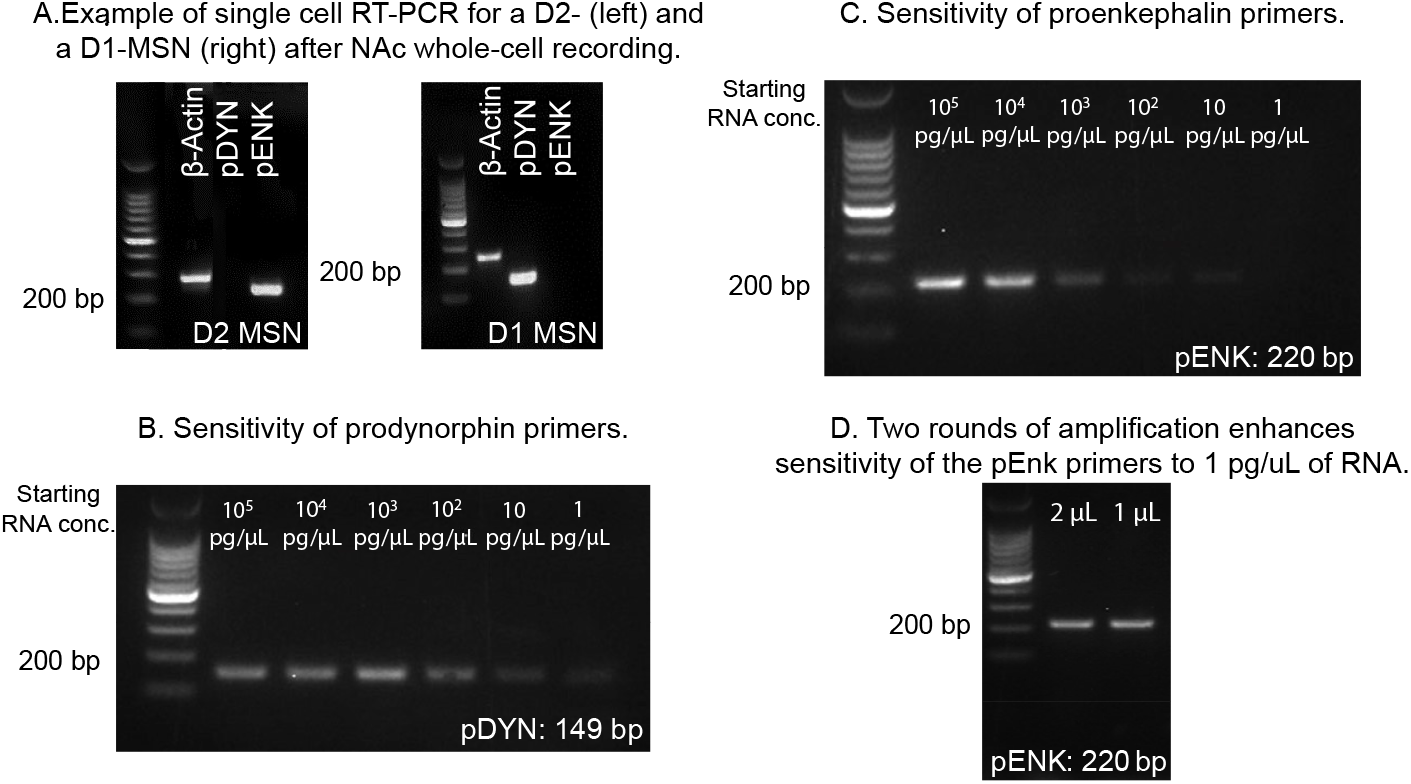
Verification of single cell RT-PCR method. A) Example of single cell RT-PCR for a D2-MSN (left) and a D1-MSN (right) after whole-cell recordings in adult rat nucleus accumbens. *β*-actin was used as a positive control. B) Serial dilution of RNA from striatal tissue showing the sensitivity of prodynorphin primers (pDYN; 149 bp). C) Serial dilution of RNA from striatal tissue showing the sensitivity of proenkephalin primers (pENK; 220 bp). D) A second round of amplification is sufficient to allow for proenkephalin detection in samples containing 1 pg/*µL* of RNA.

**FIG. 3:**
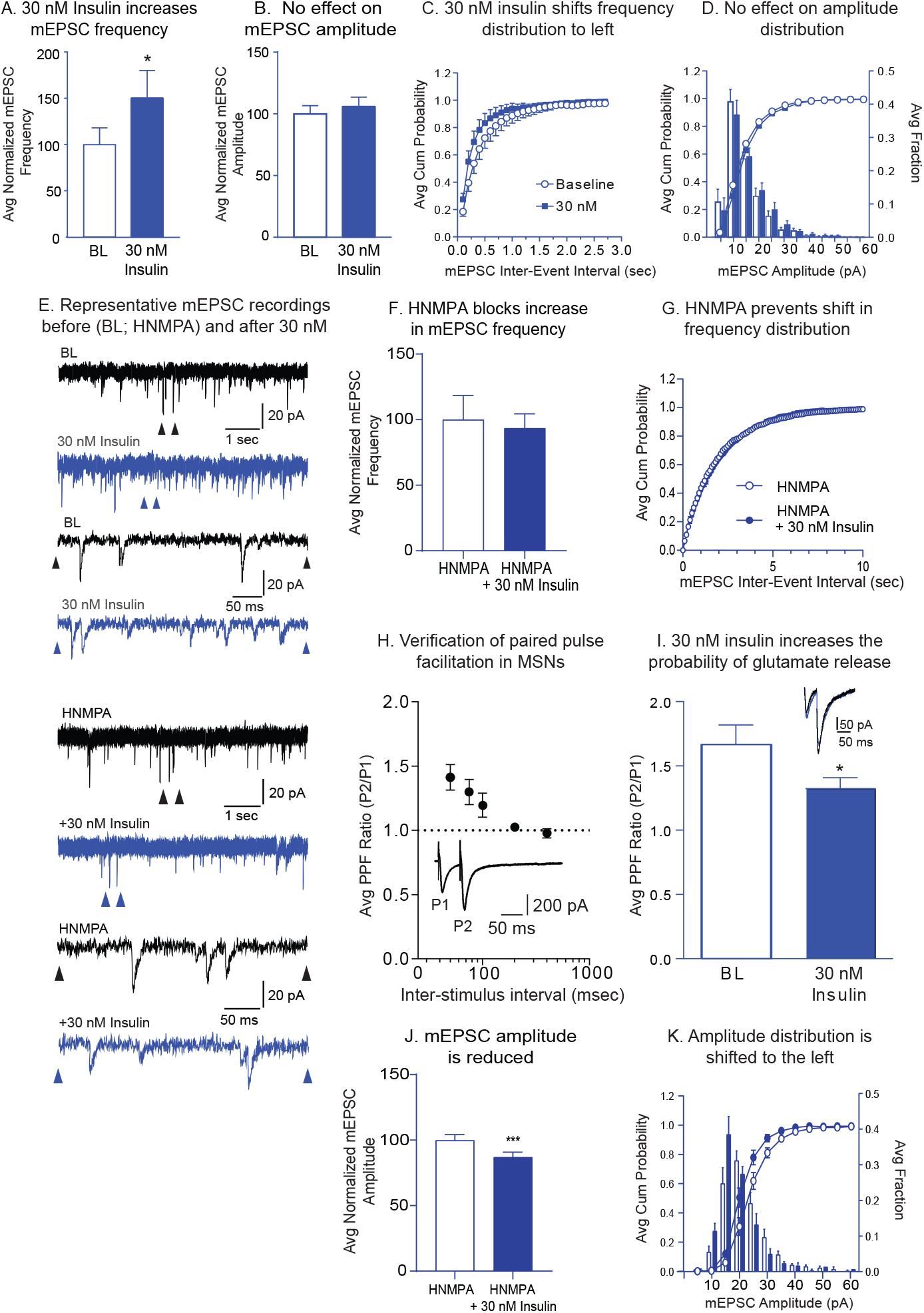
30nM insulin increases mEPSC frequency and the probability of glutamate release, but not mEPSC amplitude. A) Average mEPSC frequency before (baseline, BL) and after bath application of insulin (30nM). B) Average mEPSC amplitude before and after insulin (30nM). C) Cumulative probability distributions of mEPSC frequency before and after insulin (30nM). D) Cumulative probability distributions and histograms of mEPSC amplitude before and after insulin (30nM). E) Representative mEPSC traces before and after insulin (30nM), with and without the membrane-impermeable insulin receptor inhibitor HNMPA included in the patch pipette. Arrows in the upper traces indicate regions in which the time scale was expanded in the lower traces. F) Average mEPSC frequency before and after bath application of insulin (30nM) with membrane impermeable HNMPA included in the patch pipette. G) Cumulative probability distributions of mEPSC frequency before and after insulin (30nM) with HNMPA included in the patch pipette. H) Verification of paired-pulse facilitation in MSNs. Average PP ratio across increasing inter-pulse intervals (50-400 ms). Inset shows representative traces at a 50ms inter-pulse interval. As expected, the probability of glutamate release is relatively low in NAc medium spiny neurons, and facilitation occurs at inter-pulse intervals at or below 100 msec. I) Average PP ratio before and after insulin (30nM). Representative traces before (black) and after insulin (gray) are shown in the inset. J) Average mEPSC amplitude before and after insulin (30nM) with membrane impermeable HNMPA included in the patch pipette. K) Cumulative probability distributions and histograms of mEPSC amplitude before and after insulin (30nM) with HNMPA included in the patch pipette. Statistical differences were determined by two-tailed paired t-tests; ∗ = p < 0.05.

**FIG. 4:**
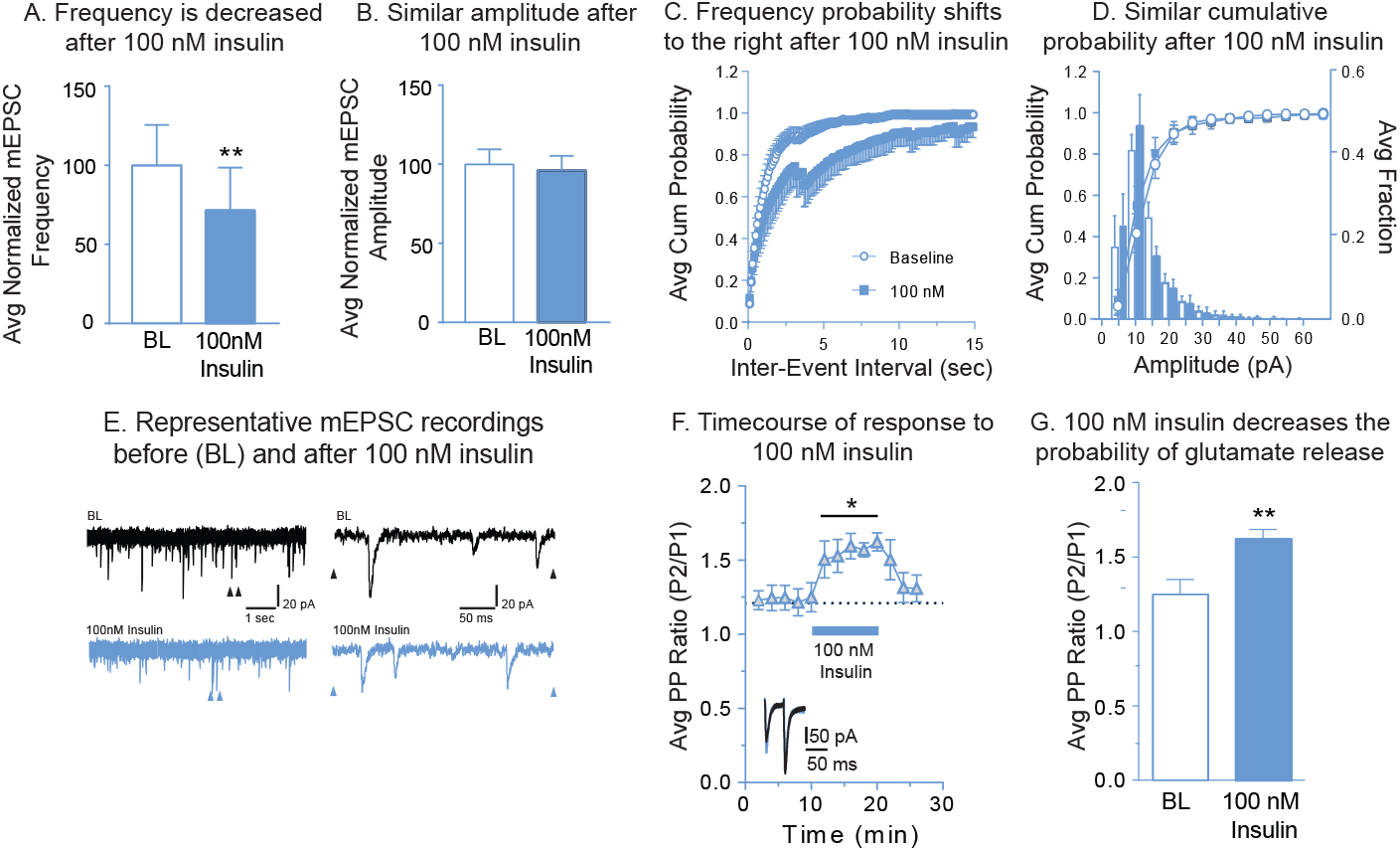
100 nM insulin reduces mEPSC frequency and the probability of glutamate release without altering mEPSC amplitude. A) Average mEPSC frequency before (baseline, BL) and after insulin (100 nM). B) Average mEPSC amplitude before and after insulin (100 nM). C) Cumulative probability distributions of mEPSC frequency before and after insulin (100 nM). D) Cumulative probability distributions and histograms of mEPSC amplitude before and after insulin (100 nM). E) Representative mEPSC traces before and after insulin (100 nM). Arrows in the left traces indicate regions in which the time scale was expanded in traces shown at the right. F) Average PP ratio before and after insulin (gray bar, 100 nM) and following insulin wash out. Representative traces before (black) and after insulin (gray) are shown in the inset. G) Average change in the PP ratio following insulin (100nM). Statistical differences were determined by two-tailed paired t-tests (A, G; ** = p < 0.01) and two-way RM ANOVA comparing baseline and treatment conditions (F; ∗ = main effect of treatment, p < 0.01).

**FIG. 5:**
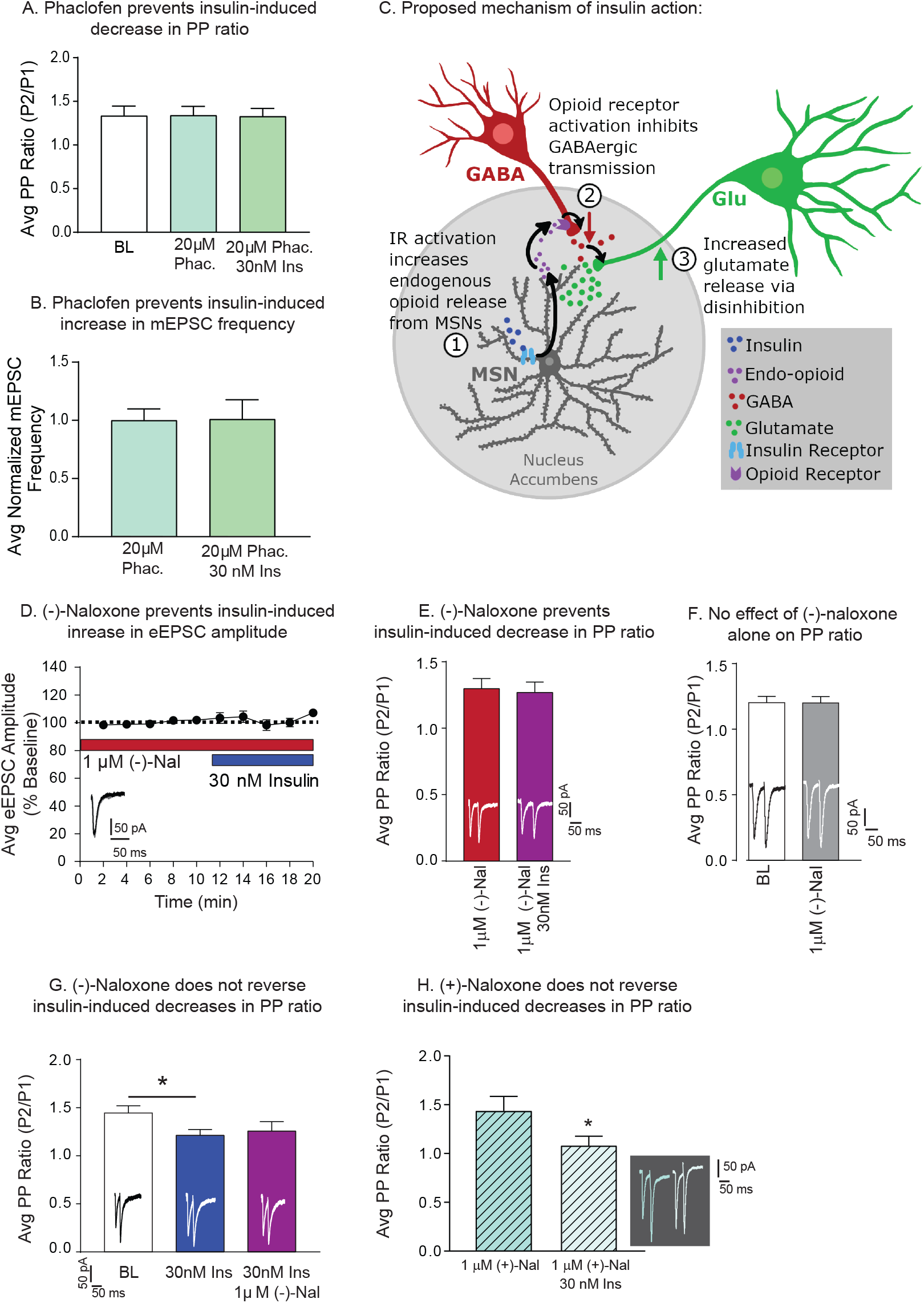
A) Average PP ratio at baseline (BL), and after insulin (30nM) in the absence and presence of the GABA_*B*_ receptor antagonist Phaclofen (20*µM*). B) Average mEPSC frequency before and after insulin (30nM) in the presence of Phaclofen (20 *µM*). C) Proposed mechanism by which activation of insulin receptors on MSNs enhances glutamate release. We propose that activation of insulin receptors on MSNs results in the release of endogenous opioids (1) that reduces GABAergic transmission (2), thereby causing disinhibition of presynaptic glutamate release (3). D) Average eEPSC amplitude in the presence of the opioid receptor antagonist (-)-naloxone (1 *µM*) before and after bath application of insulin (30 nM). E) Average PP ratio in the presence of (-)-naloxone (1 *µM*) before and after bath application of insulin (30 nM). F) Average PP ratio before and after bath application of (-)-naloxone (1 *µM*) alone. G) Average PP ratio at baseline (BL), and after insulin (30 nM) in the absence or presence of (-)-naloxone (1 *µM*). H) Average PP ratio in the presence of (+)-naloxone (1 *µM*) before and after bath application of insulin (30 nM). Example traces are shown within each panel. Statistical differences were determined by two-way RM ANOVA comparing baseline and treatment conditions (D), two-tailed paired t-tests (C, E, F, H), and one-way ANOVA followed by Sidaks multiple comparisons post-test (B,G); ∗ = p < 0.05.

Of the 72 cells collected, 13 cells could not be classified because bands were not visible (likely due to low starting RNA content). Of the cells identified, 41% were D2-MSNs and 47% were D1-MSNs, consistent with the literature (Sun et al., 2008). The remaining 12% were positive for both pENK and pDYN. This dual expression could be due to contamination from other cells as the pipette was removed from the slice. While this single cell RT-PCR method can distinguish D1- and D2-MSNs, statistical comparisons were not possible due to low N within a given measure in the current study (in part due to low RNA starting content, or inability to collect cell contents before the end of the recording). Thus, although no overt differences between D1- and D2-MSNs in response to insulin were apparent (data not shown), future studies utilizing transgenic rats specifically designed to identify MSN sub-populations can be used to further investigate potential cell specific effects (Pettibone et al., 2018). However, it is worthwhile to note that the variance of insulins effects on transmission were low overall, suggesting that effects are consistent across the MSNs we recorded from. This could be due to cross-talk between D1- and D2-MSNs, and/or afferent glutamatergic input to both cell populations.

### D. Insulin-induced changes in excitatory transmission are mediated by alterations in presynaptic glutamate release

In cultured hippocampal cells, insulin enhances endocytosis of GluA2-containing AMPARs, while also increasing exocytosis GluA1-containing AMPARs (Beattie et al., 2000; Man et al., 2000; Passafaro et al., 2001; Huang et al., 2004). In addition, more recent studies have found that insulin receptor activation in the ventral tegmental area (VTA) reduces presynaptic glutamate release (Labouebe et al., 2013; Liu et al., 2013). Thus, we next examined the effect of 30 or 100 nM insulin on miniature EPSC (mEPSC) amplitude and frequency, and on paired pulse facilitation (Figures 3, 4). 30 nM Insulin increased mEPSC frequency (Figure 3A; two-tailed paired t-test *t*_5_=3.45, p=0.02; N=6,5) without altering mEPSC amplitude (Figure 3B; two-tailed paired t-test: *t*_5_=1.66, p=0.16). In addition, the frequency cumulative probability distribution was shifted to the left compared to baseline (Figure 3C), with no change in the amplitude cumulative probability distribution (Figure 3D). Furthermore, including HNMPA in the recording pipette blocked this insulin-induced increase in mEPSC frequency (Figure 3F; two-tailed paired t-test: *t*_5_=0.465, p=0.66, N=6,3) and prevented the leftward shift in the frequency cumulative probability distribution (Figure 3G). These data suggest insulin-induced increases in excitatory transmission are mediated by enhanced glutamate release. To confirm these results we recorded from additional neurons using a paired pulse facilitation procedure. We first verified the paired pulse method in our hands by measuring eEPSCs across a range of inter-pulse intervals. As expected facilitation occurred at or below an inter-pulse interval of 100 ms (Figure 3 H) and the probability of glutamate release on NAc MSNs was relatively low (Kasanetz and Manzoni, 2009). Consistent with effects on mEPSC frequency, 30 nM insulin decreased the paired pulse ratio (Figure 3G; two-tailed paired t-test *t*_4_=3.32, p=0.03; N=5,4), indicative of an increase the probability of glutamate release. Taken with data above this suggests that activation of insulin receptors increases excitatory transmission in the NAc core by enhancing glutamate release onto MSNs. In addition, when HNMPA was included in the recording pipette and 30 nM insulin was applied, a decrease in mEPSC amplitude was found (Figure 3J; two-tailed paired t-test: *t*_5_=7.75, p=0.006, N=6, 3) and the cumulative probability distribution of mEPSC amplitudes were shifted to the left (Figure 3K). This suggests additional effects of insulin on post-synaptic transmission that are not mediated by insulin receptor activation.

Effects of 100 nM insulin on mEPSC amplitude and frequency (Figure 4A-E), and paired pulse facilitation (Figure 4F, G) were also examined. We found that the frequency of mEPSCs was significantly reduced by 100 nM insulin (Figure 4A; two-tailed paired t-test *t*_6_=3.90, p=0.008; N=7,6), without altering mEPSC amplitude (Figure 4B; two-tailed paired t-test *t*_6_=0.94, p=0.38). In addition, the frequency cumulative probability distribution was shifted to the right (Figure 4C), with no change in the amplitude cumulative probability distribution (Figure 4D). Consistent with reductions in mEPSC frequency, 100 nM insulin significantly increased the paired pulse ratio (Figure 4F; two-way RM ANOVA main effect 100 nM: *F*_(1,6)_=21.41, p=0.003; Figure 4G; two-tailed paired t-test *t*_6_=4.67, p=0.003; N=7,5), indicating a reduction in the probability of glutamate release following 100nM insulin. Thus, the data suggest that activation of IGFRs by insulin reduces excitatory transmission in the NAc core by decreasing glutamate release onto MSNs.

### E. Insulin-induced increases in excitatory transmission are due to opioid receptor-mediated disinhibition

Endogenous concentrations of insulin in the brain are thought to range from 30-50 nM (Havrankova et al., 1978; Schulingkamp et al., 2000). Therefore, we focused our studies of underlying mechanisms on insulin-receptor mediated increases in presynaptic glutamate release following 30 nM insulin. Because blockade of insulin receptors within the recorded MSN was sufficient to prevent increases in excitatory transmission following 30nM insulin (Figure 1D, 3F), we reasoned that presynaptic effects are likely mediated by a neuromodulator released by MSNs, such as GABA or endogenous opioids. Given that both of these transmitters are inhibitory, it is unlikely that effects of insulin are due to direct effects on presynaptic glutamatergic terminals, as activation of GABA or opioid receptors on glutamatergic terminals reduces presynaptic glutamate release, not enhances it (Nisenbaum et al., 1993; Hjelmstad and Fields, 2003). Therefore, we hypothesized that effects may be due to disinhibition of inhibitory inputs onto glutamatergic terminals (as GABA_*A*_ but not GABA_*B*_ receptors were blocked during our recordings). This idea is also based in part on the ability of GABA_*B*_, but not GABA_*A*_, receptors on glutamatergic terminals to decrease glutamate release in the striatum (Nisenbaum et al., 1993). Consistent with our hypothesis, addition of the GABA_*B*_ receptor antagonist phaclofen (20 *µM*) to the bath was sufficient to prevent insulin-induced increases in glutamate release measured using both PPF (Figure 5A; one-way RM ANOVA no effect of 30 nM insulin: *F*_(2,8)_=0.074, p=0.93, N=5,3) and mEPSC frequency (Figure 5B; two-tailed paired t-test: *t*_5_=0.082, p=0.94, N=5,3). Thus, insulin-induced increases in excitatory transmission appear to rely on disinhibition, rather than direct enhancement of glutamate release. Importantly, the concentration of phaclofen used did not affect basal transmission, suggesting that we did not simply enhance excitatory transmission to a ceiling (not shown).

Opioid receptors are found on GABAergic terminals in the NAc (Pickel et al., 2004), and activation of opioid receptors causes disinhibition in the VTA and hippocampus by reducing GABAergic transmission (Capogna et al., 1993; Hjelmstad et al., 2013). Thus, we speculated that insulin may trigger endogenous opioid release, which then activates opioid receptors on GABAergic terminals within the NAc to disinhibit presynaptic glutamate release (see schematic Figure 5A; note that recordings were done in coronal slices which contain cell bodies of cells intrinsic to the NAc and terminals, but not cell bodies, from regions that project to the NAc). Therefore, we next determined whether application of the opioid receptor antagonist (-)-naloxone (1 *µM*, Chieng and Christie, 1994) would prevent insulin-induced increases in excitatory transmission.

Bath application of (-)-naloxone prior to 30nM insulin prevented insulin-induced increases in eEPSC amplitude (Figure 5D; two-way RM ANOVA no effect of 30 nM insulin: *F*_(1,5)_=0.82, p=0.41; N=6,3), and insulin-induced reductions in paired pulse ratio (Figure 5E; two-tailed paired t-test, *t*_6_=0.43, p=0.68; N=7,4). In addition, (-)-naloxone alone did not alter the probability of glutamate release in the absence of insulin (Figure 5F; twotailed paired t-test *t*_9_=0.53, p=0.53; N=10,5). This suggest that opioid receptor activation is secondary to insulin receptor activation. Given these results, its logical to suspect that enhancing basal opioid tone could partially occlude insulins effects. In an attempt to address this possibility, we utilized the peptidase inhibitors bestatin (10 *µM*) and thiorphan (1 *µM*), which can prevent the degradation of endogenous opioids (Birdsong et al., 2019). Bath application of these peptidase inhibitors decreased eEPSC amplitude from baseline, with no further changes observed when 30nM insulin was applied (data not shown; two-way RM ANOVA main effect of treatment: *F*_(2,6)_=33.2, p¡0.001; N=4,3; Sidaks multiple comparisons post-test: BL vs. peptidase inhibitors, p=0.003; peptidase inhibitors vs. 30nM, p=0.31). The large reduction in eEPSC amplitude observed when the peptidase inhibitors were bath applied is consistent with a generalized inhibitory effect of increasing opioid tone, but complicates the interpretation of subsequent insulin application. Thus although the effect of insulin was occluded in these recording conditions, consistent with data above, this could merely be due to the overall inhibition caused by enhancing opioid tone. It is worthwhile to note that assessment of basal striatal opioid tone has been difficult (see Birdsong and Williams, 2020 for discussion of this point). However, reductions in eEPSC amplitude induced by the peptidase inhibitors provide some evidence for accumulation of opioids within our acute slices. This is consistent with one previous report in the dorsal striatum (Atwood et al., 2014).

Application of (-)-naloxone after 30 nM insulin (10 min) was not sufficient to reverse insulin-induced decreases in PPF (Figure 5G; two-way RM ANOVA main effect of treatment *F*_(2,6)_=6.61, p=0.03; N=4,3; Sidaks multiple comparisons post-test BL vs. 30nM, p=0.04). This is consistent with the absence of effects of (-)- naloxone alone and suggests that once opioid receptor signaling is triggered, subsequent opioid receptor blockade cannot overcome ongoing signaling. Finally, to more conclusively test the role of opioid receptor activation, we bath applied (+)-naloxone (1 *µM*) prior to 30 nM insulin. (+)-Naloxone is the structural enantiomer of (-)-naloxone, but does not have any action at opioid receptors (Iijima et al., 1978). Consistent with the data above, (+)-naloxone did not prevent insulin-induced increases in glutamate release measured by paired pulse facilitation (Figure 5H; two-tailed paired t-test *t*_7_=2.55, p=0.04; N=8,5). Taken together, antagonist studies using (-) and (+)-naloxone show that insulin-induced increases in presynaptic glutamate release require opioid receptor activation.

### F. Diet-induced obesity blunts insulin-receptor mediated increases in excitatory transmission and reduces NAc insulin receptor surface expression

Circulating insulin reaches the NAc and diet-induced obesity is accompanied by chronic elevations in circulating insulin (Woods et al., 2016). In addition, obesity has been associated with a reduction of cognitive-enhancing effects of intra-nasal insulin in humans see (see (Kullmann et al., 2016) for review) and impairments in hippocampal glutamatergic plasticity (Fadel and Reagan, 2016). Therefore, we predicted that chronic elevations in endogenous insulin produced by high-fat diet-induced obesity may blunt insulin’s ability to enhance NAc excitatory transmission. For this set of studies, adult male rats were given free access to 60 % high-fat diet in the home cage for a total of 8 weeks, while controls had free access to standard lab chow (13% fat). As expected, high-fat diet resulted in significant increases in fasted plasma insulin levels (Figure 6A; two-tailed unpaired t-test *t*_26_=3.65, p=0.001; chow N=13, high-fat N=15) and fat mass compared to controls (Figure 6B; two-tailed unpaired t-test *t*_26_=6.82, p<0.0001). We next examined the effect of bath application of 30 nM and 100 nM on eEPSC amplitude in slices from high-fat diet and control rats. Similar to results above, 30 nM insulin increased eEPSC amplitude, whereas 100 nM insulin decreased it in recordings from controls (Figure 6C circles; two-way RM ANOVA main effect insulin: *F*_(2,8)_=5.17, p=0.04; N=5,3). In contrast, in MSNs from high-fat rats 30 nM insulin did not significantly alter eEPSC amplitude, while significant decreases induced by 100 nM insulin persisted (Figure 6C squares; two-way RM ANOVA main effect insulin: *F*_(2,10)_=8.74, p=0.01; BL vs. 30 nM; main effect of insulin: *F*_(1,5)_=2.86, p=0.15; BL vs. 100 nM; main effect of insulin: *F*_(1,5)_=8.58, p=0.03; N=6,5). These data are consistent with the idea that physiological shifts in circulating insulin secondary to diet-induced obesity impact neural insulin sensitivity (see also, Ferrario and Reagan, 2017).

**FIG. 6:**
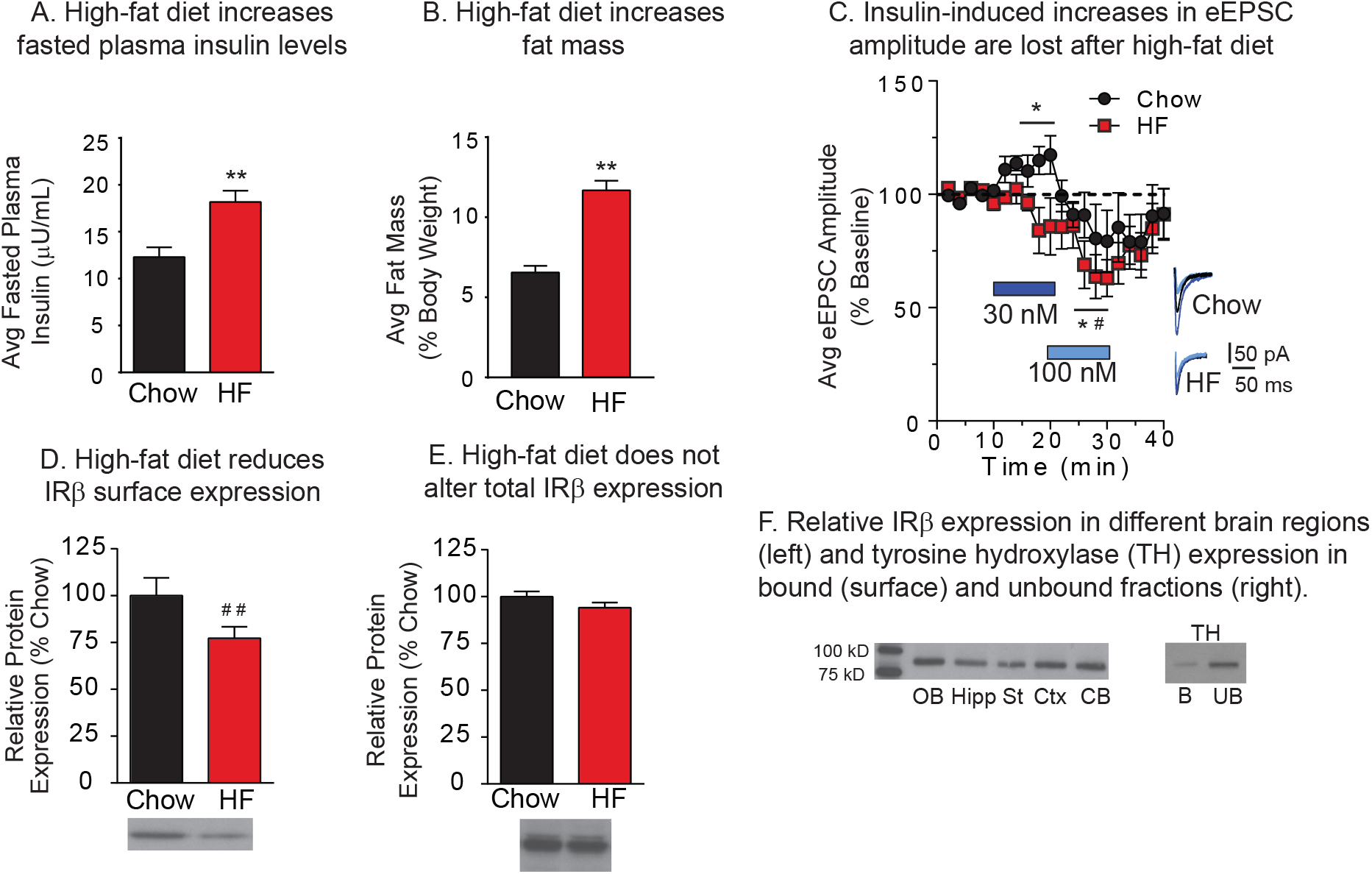
High-fat diet-induced obesity results in a loss of insulin-induced increases in excitatory transmission and a reduction in NAc insulin receptor *β* (IR*β*) surface expression. Average concentration of fasted plasma insulin (A) and fat mass (B) in chow and high-fat diet groups. C) Average eEPSC amplitude following bath application of increasing concentrations of insulin (gray bars) and following insulin wash out in MSNs from chow (open circles) and high-fat groups (closed squares). Representative traces for each group before (black) and after (gray) each insulin concentration are shown at the right. D) Average NAc IR*β* surface expression in high-fat and chow fed groups. E) Total NAc IR*β* expression in high-fat and chow fed groups. Representative blot images are shown below each graph. F) Relative abundance of IR*β*expression in the olfactory bulb (OB), hippocampus (Hipp) striatum (St), cortex (Ctx) and cerebellum (CB). Total protein in each lane did not differ between groups (not shown). G) Immunoblot for tyrosine hydroxylase (TH) in the bound (B) and unbound (UB) fractions. Consistent with its intracellular localization, TH protein levels were nearly absent in the bound (surface) fraction. Statistical differences were determined by two-tailed unpaired t-tests (A, B, D & E; ∗∗ : *p* < 0.001, ##: *p* = 0.05), and two-way RM ANOVA comparing baseline and treatment conditions (C; * = chow group, main effect of treatment, # = high-fat group, main effect of treatment, *p* < 0.05).

One potential explanation for the loss of insulin-induced increases in excitatory transmission is a reduction in NAc insulin receptor expression. Therefore, we next determined the effect of high-fat diet on surface expression of the *β* subunit (IR*β*) of the insulin receptor using established biotinylation and pull down procedures (Ferrario et al., 2011). Relative expression in whole cell lysates across brain regions was used to validate the IR*β* antibody (Figure 6F). Consistent with the literature, IR*β* was most abundant in cerebellum and olfactory bulb compared to other regions (Ponceau S staining was used to verify equal protein loading per lane, data not shown). In addition, the intracellular protein TH was apparent in the unbound, but not the bound, fraction (Figure 6G), verifying enrichment of surface proteins. We found a 21.7% (*±* 7.6%)*reductioninNAcIRβ* surface expression in high-fat vs. chow fed groups (Figure 6D; two-tailed unpaired t-test *t*_18_=1.75, p=0.05) without any changes in total IR*β* (Figure 6E). This suggests that the loss of insulin-induced increases in eEPSC amplitude in the high-fat diet group may be due to reductions in NAc insulin receptor expression.

## IV. DISCUSSION

### A. Insulin receptor and IGFR activation have opposing effects on NAc excitatory transmission

We found bi-directional effects of insulin on NAc excitatory transmission, with relatively low concentrations of insulin increasing the eEPSC amplitude, and concentrations over 100 nM decreasing it (Figure 1A, B). Using a combination of antagonists and blockers, we show that increases in excitatory transmission are mediated by insulin receptors whereas decreases are mediated by IGFRs. Furthermore, insulin receptor-mediated effects were attributable to activation of insulin receptors on MSNs, as inclusion of a membrane impermeable insulin receptor blocker in the recording pipette was sufficient to completely prevent insulin induced increases in excitatory transmission. Interestingly, the magnitude of the insulin-induced increase in eEPSC amplitude was similar in the presence or absence of an IGFR antagonist (∼ 25-30%). This suggests that 30 nM insulin only activates insulin receptors whereas higher insulin concentrations are required to recruit IGFR activation. Indeed, 50nM insulin did not alter excitatory transmission, presumably because the sum of enhancing (insulin receptormediated) and reducing (IGFR-mediated) transmission were off-setting. These results suggest that the net effect of insulin on excitatory transmission in vivo may be strongly influenced by local insulin concentration (see also, effects of high-fat diet below).

We next determined whether insulin-induced changes in excitatory transmission are due to alterations of pre- or postsynaptic function. Previous studies in hippocampus have found that insulin can enhanced AMPAR endocytosis and exocytosis, see results and (Ferrario and Reagan, 2017) for review. However, we did not find any evidence for changes in postsynaptic AMPAR transmission. Instead, application of 30 nM insulin increased mEPSC frequency and the probability of glutamate release without altering mEPSC amplitude (Figure 3). To our knowledge, this is the first time insulin has been found to enhance excitatory transmission via a presynaptic mechanism. In contrast, in VTA, insulin receptor activation produces rapid and persistent reductions in presynaptic glutamate release (Labouebe et al., 2013; Liu et al., 2013). Thus, effects of insulin are region specific, although it should be noted that previous studies were conducted in cultured neurons or juvenile mice, while studies here are in adult rats. Finally, IGFR-mediated reductions in excitatory transmission were also due to effects on presynaptic glutamate release (Figure 4). This is consistent with the ability of IGFR activation to inhibit L-type calcium channel activity which mediates presynaptic glutamate release (Subramanian et al., 2013; Sanchez et al., 2014), and with the ability of IGFR activation to suppress spontaneous excitatory transmission in hippocampus (Gazit et al., 2016).

### B. Insulin-induced increases in excitatory transmission are due to opioid receptor-mediated disinhibition

Blockade of insulin receptor signaling within the recorded MSN was sufficient to prevent insulin-induced increases in excitatory transmission (Figure 1D), suggesting a mechanism involving feedback from MSNs to presynaptic glutamatergic terminals. Indeed, our data support a previously unidentified mechanism whereby insulin produces disinhibition mediated by opioid receptor activation (Figure 5A). As described above, recordings were made in the presence of a GABA_*A*_ antagonist, thus ionotropic inhibition cannot contribute. Furthermore, addition of a GABA_*B*_ antagonist prevented insulin-induced increases in release probability and mEPSC frequency (Figure 5A,B). While removing all GABA transmission is quite a hammer, this nonetheless provides additional support for disinhibition. Furthermore, addition of the opioid receptor antagonist (-)-naloxone was sufficient to prevent insulin-induced increases in excitatory transmission measured by mEPSC frequency, paired pulse ratio, and eEPSC amplitude (Figure 5A-E). The role of opioid receptors was further supported by the inability of (+)-naloxone (which does not have any action at opioid receptors (Iijima et al., 1978)) to prevent insulin-induced increases in glutamate release (Figure 5H). Furthermore, naloxone alone was not sufficient to alter glutamate release (Figure 5F), suggesting that there is not opioid-mediated tonic inhibition of presynaptic glutamate. However, activation of insulin receptors on MSNs could acutely trigger endogenous opioid release, and thus opioid tone is not necessarily required for the observed effects of insulin.

Although few functional studies have examined the regulation of NAc glutamate release by endogenous opioids, this mechanism is consistent with anatomical and physiological data. Specifically, Mu opioid receptors are located on presynaptic GABAergic terminals within the NAc (Svingos et al., 1997; Pickel et al., 2004), and Mu opioid receptor activation reduces GABA release in the hippocampus and sub-thalamic nucleus (Lambert et al., 1991; Xie et al., 1992; Capogna et al., 1993; Lupica, 1995; Shen and Johnson, 2002). In addition, GABAB receptors are located on glutamatergic terminals within the striatum where they inhibit excitatory transmission (Nisenbaum et al., 1993). Thus, it is feasible for endogenous opioids to produce the disinhibition observed here (Shen and Johnson, 2002; Banghart et al., 2015; Tejeda et al., 2017). Finally, naloxone is a non-selective opioid receptor antagonist. Kappa opioid receptors are located on terminals of excitatory and inhibitory synapses within the NAc (Svingos et al., 1999; Meshul and McGinty, 2000; Tejeda et al., 2017), and on dopamine afferents in the NAc (Spanagel et al., 1992), whereas delta opioid receptors are preferentially expressed on cholinergic interneurons within the NAc (Le Merrer et al., 2009; Bertran-Gonzalez et al., 2013; Castro and Bruchas, 2019). Thus, in addition to potential roles for Mu opioid receptors discussed above, effects observed here could be mediated by one, or a combination of different opioid receptors. Future studies are needed to determine the receptor population(s) involved.

### C. Potential contribution of dopamine

Insulin modulates dopamine systems at the level of VTA dopamine neurons (Mebel et al., 2012; Labouebe et al., 2013; Liu et al., 2013) and at dopamine terminals within the NAc (Stouffer et al., 2015; Naef et al., 2018). Within VTA, relatively high concentrations of insulin (100-500 nM) reduce presynaptic glutamate release onto VTA dopamine neurons (Labouebe et al., 2013; Liu et al., 2013), while within the NAc 30 nM, but not 100 nM, insulin enhances dopamine release by increasing cholinergic interneuron firing (Stouffer et al., 2015). While it is tempting to suggest that insulin-induced increases in NAc dopamine or interneuron activity may contribute to increases in excitatory transmission found here, two lines of evidence argue against this. First, insulin-induced increases in NAc dopamine release do not occur until ∼ 20 min following insulin application, whereas the onset of effects here were immediate and maximal within ∼ 5 min. Second, blockade of insulin receptors only within the recorded neuron here completely prevented insulin-induced increases in excitatory transmission (Figure 1D, 3F). Given that this manipulation would not prevent activation of insulin receptors on other cells within the slice, it seems unlikely that effects here are secondary to insulin-induced changes in dopamine (see also (Ferrario and Reagan, 2017) for additional discussion).

### D. Loss of insulin-receptor mediated effects following obesity

When effects of high-fat diet were examined, we found a loss of insulin receptor-mediated increases in excitatory transmission, but a maintenance of IGFR-mediated decreases (Figure 6C). Although slight trends were seen for reduced transmission following 30nM insulin in the high-fat group, the p value indicated a low probability of a true effect. The loss of insulin-induced increases is likely due, at least in part, to a reduction in NAc insulin receptor expression, as surface expression of IR*β* was reduced following high-fat diet (Figure 6D). These data are consistent with the development of insulin resistance in the face of chronic elevations in circulating insulin resulting from obesity, and with impairments in hippocampal glutamatergic plasticity induced by insulin resistance (Fadel and Reagan, 2016). Although we cannot rule out the contribution of differences in basal insulin tone between chow and high-fat groups, these data nonetheless demonstrate that physiologically relevant increases in circulating insulin are accompanied by reductions in insulin receptor-mediated effects on NAc excitatory transmission.

### E. Summary and future directions

Studies above provide the first insights into how insulin influences NAc excitatory transmission. Insulins effects and the underlying mechanism in the NAc differ dramatically from those found in other brain regions and at earlier stages in neural development. Furthermore, the involvement of opioid receptor activation in insulin-receptor mediated increases in glutamate release found here is consistent with the ability of opioids in the NAc to enhance food intake and hedonic responses to palatable foods (Zhang and Kelley, 2000; Katsuura and Taha, 2013; Richard et al., 2013; Castro and Bruchas, 2019), and with conditioned increases in insulin in anticipation of palatable food consumption (Teff, 2011). Thus, results here have broad implications for the regulation of food-seeking and eating behavior by insulin, as well as motivation for non-essential reinforcers like cocaine (Naef et al., 2018) that also relies on NAc excitatory transmission. Finally, the NAc receives inhibitory input from GABAergic neurons in the VTA (Van Bockstaele and Pickel, 1995), in addition to local collateralization of MSNs and aspiny GABAergic interneurons (Smith and Bolam, 1990; Kawaguchi, 1993; Planert et al., 2010). Thus, in addition to identifying the opioid receptors involved, it will be important for future studies to determine whether disinhibition produced by insulin is selective to different sources of GABA within the NAc.

## Acknowledgments

Artwork in Fig 5C by Victoria Macht, PhD. This work was supported by NIDDK R01DK106188, R01DK115526, and the Brain and Behavior Research Foundation N018940 awards to CRF; TLF and MFO were supported by NIDA T32DA007268 MFO was supported by NIDDK FDK112627A; YAC was supported by 3R01DK106188-02-S1 and 1F99NS108549-01. The authors declare no competing financial interests. Studies utilized the Chemistry Core of the Michigan Diabetes Research and Training Center funded by DK020572, and the University of Michigan Animal Phenotyping Core supported by P30 grants DK020572 (MDRC) and DK089503 (MNORC). We thank Drs. Marina Wolf, Larry Reagan, William Birdsong, and Travis Brown for helpful conversations and feedback on these studies, Dr. Garret Stuber for helpful comments, and Dr. Kenner C. Rice (Drug Design and Synthesis Section, NIDA IRP) for the gift of (+)-naloxone.

## Author Contributions

TLF, MFO conducted experiments, and contributed to experimental design, data analysis and wrote the paper; AMN and YAC conducted experiments, data analysis, and contributed to writing of the paper; ZSR conducted pull down experiments and wrote corresponding results; CRF contributed to experimental design, conducted analysis, and wrote the paper.

